## Supplementary Figures 1 to 4 for "Lifecycle of a predatory bacterium vampirizing its prey through the cell envelope and S-layer"

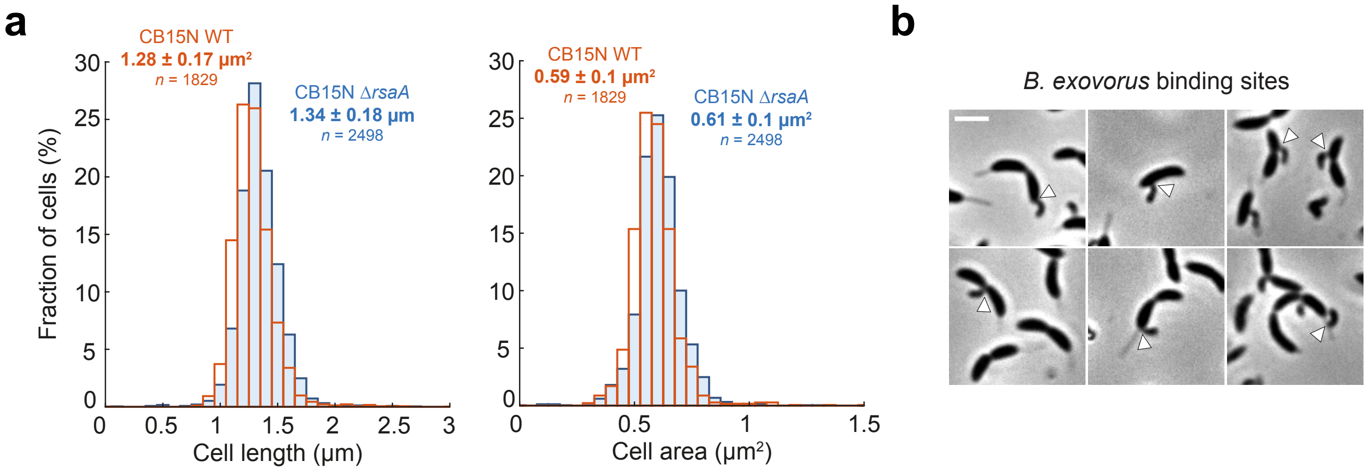


**Supplementary Fig. 1 | The presence of the S-layer does not dictate the predator binding site or the size of newborn predators. a**, Histograms representing the indicated cellular dimensions of the selected *B. exovorus* predators upon overnight predation of either the wild-type (CB15N WT, orange) or the Δ*rsaA* *C. crescentus* (CB15N Δ*rsaA*, blue) as a prey. Quantification based on the predator cells depicted on **Fig. 1a** and **Fig. 2a**. Values of the mean and the standard deviation, and the number of analyzed predator cells (*n*) are indicated on the graphs for each condition. **b**, Representative phase contrast images of *B. exovorus* cells attached to the wild-type *C. crescentus* upon 15 min of co-incubation. White arrowheads highlight predator binding sites. All selected binding sites result in predator growth. Scale bar, 2 µm. Related to **Fig. 3**.

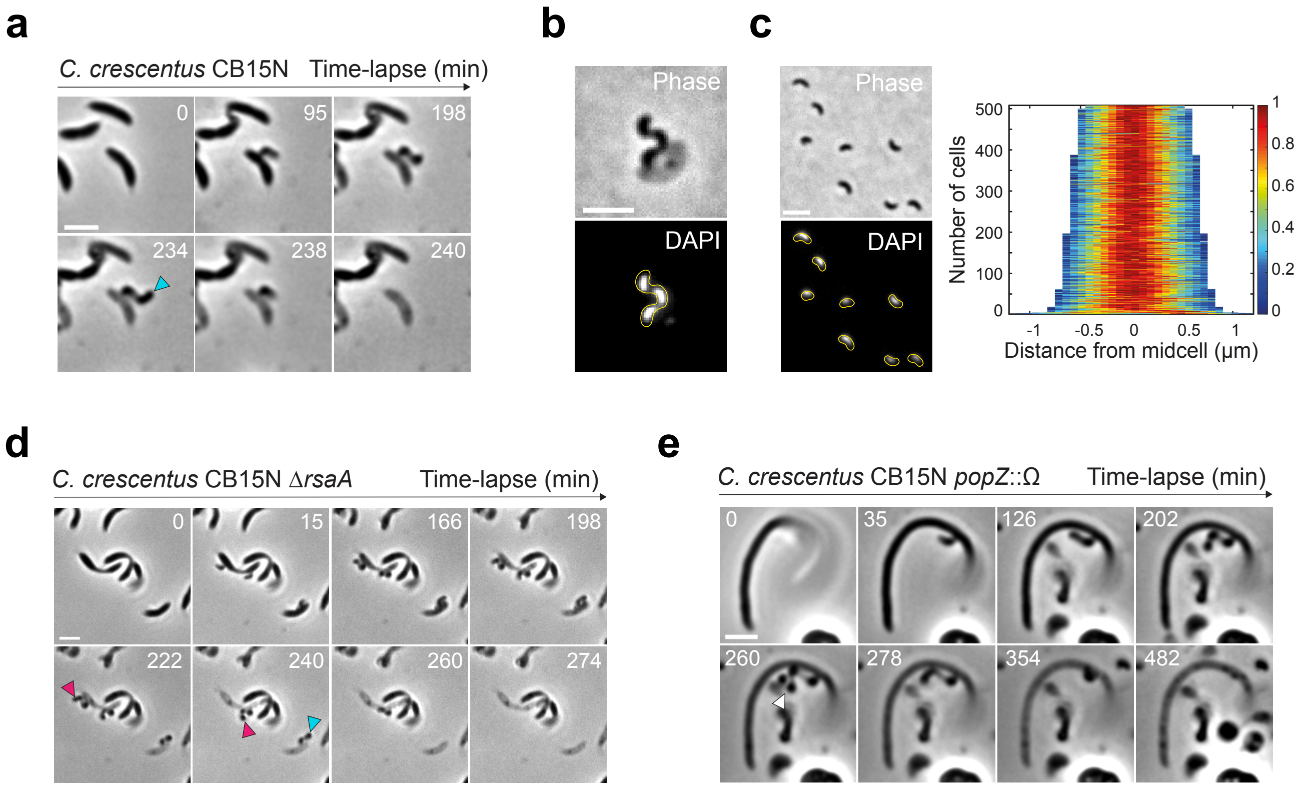


**Supplementary Fig. 2 | The predator offspring is unaffected by the presence of the S-layer or the prey cell size. a**, Representative time-lapse phase contrast microscopy images of the *B. exovorus* growth using the wild-type *C. crescentus* strain as a prey. The cyan arrowhead shows the production of 2 progenies. Scale bar, 2 µm. **b,** Representative microscopy image of a *B. exovorus* cell in the final predation stage on the surface of a ghost wild-type *C. crescentus* cell, showing future triplet progenies stained with DAPI. Top, phase contrast; bottom, DAPI fluorescence signal and *B. exovorus* cell outline drawn manually based on the phase contrast image. **c,** Left: Representative microscopy image of attack-phase *B. exovorus* upon overnight predation on wild-type *C. crescentus* and staining with DAPI. Top, phase contrast; bottom, DAPI fluorescence signal and cell outlines obtained with Oufti. Right: Demograph of the DAPI signal obtained from the same population of cells as on the left. The heatmap represents relative fluorescence intensities. Cells are sorted by length. **d-e**. Representative time-lapse phase contrast microscopy images of *B. exovorus* growth using the Δ*rsaA* (**c**) or the *popZ*::Ω (**d**) *C. crescentus* strains as a prey. Cyan, magenta, and white arrowheads show the production of 2, 3 or 4 progenies, respectively. Scale bars, 2 µm. Related to **Fig. 3**.

**Supplementary Fig. 2**

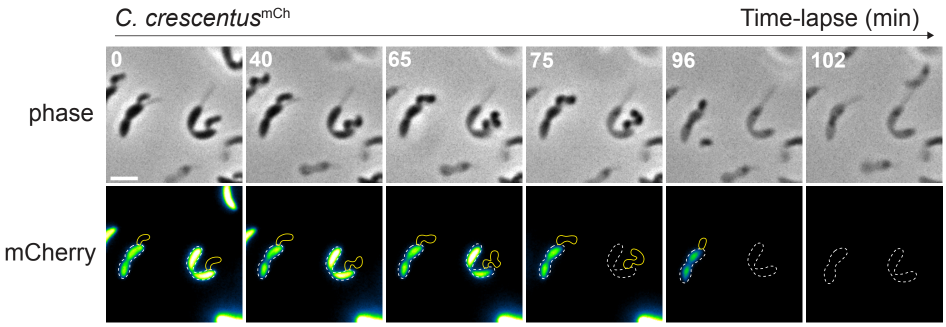


**Supplementary Fig. 3 | *In situ* digestion of the prey's proteinaceous content by *B. exovorus*.** The mCherry fluorescent signal is used as a reporter of the proteinaceous cytoplasmic content. Representative time-lapse microscopy of the mCherry-producing *C. crescentus* (*C. crescentus*$^{mCh}$) predated by *B. exovorus*. *B. exovorus* cell outlines (yellow) and *C. crescentus* prey cell outlines (dashed white) were drawn manually based on the phase contrast images. The fluorescence signal was false colored with the GreenFireBlue colormap in Fiji to display changes in fluorescence intensity. Scale bar, 2 µm. Related to **Fig. 4**.

**Supplementary Fig. 3**

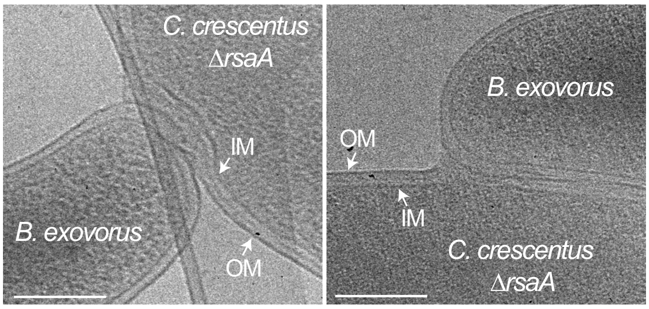


**Supplementary Fig. 4 | Cryo-EM imaging of predator:prey junction using the Δ*rsaA* C. crescentus as a prey.** Representative cryo-EM images of *B. exovorus* attached to the Δ*rsaA C. crescentus* cell surface. OM, outer membrane; IM, inner membrane. Scale bar, 0.2 μm. Related to **Fig. 5**.
