## Supplementary Table 1 for "Lifecycle of a predatory bacterium vampirizing its prey through the cell envelope and S-layer"

### Supplementary Table. 1. Bacterial strains used in this study.

| *Bdellovibrio exovorus JSS* | | |
|---|---|---|
| **Strains** | **Description** | **Source** |
| **GL1867** | JSS | ATCC BAA-2330 |
| ***Caulobacter crescentus*** | | |
| **Strains** | **Description** | **Source** |
| **GL14** | Wild-type *C. crescentus* CB15N, $Nal^R$ | Lab collection |
| **GL1866** | *C. crescentus* CB15N Δ*rsaA* (S-layer deficient mutant), $Nal^R$ | Kind gift from Régis Hallez, UNamur |
| **GL2288** | CB15N cc1959::pHU1-yfp, $Nal^R$, $Gent^R$ | Kind gift from Christine Jacob-Wagner, Stanford University |
| **GL2339** | CB15N xylX::pXbiofab-mCherry, $Nal^R$, $Kan^R$ | This study |
| **Others** | | |
| **GL1891** | *Agrobacterium tumefaciens* | Kind gift from Xavier De Bolle, UNamur |
| **GL1892** | *Sinorhizobium meliloti* | Kind gift from Xavier De Bolle, UNamur |
| **GL1893** | *Ochrobactrum anthropii* | Kind gift from Xavier De Bolle, UNAmur |
| **GL2073** | *Asticcacaulis excentricus* | Kind gift from Yves Brun, UDEM |
| **GL2076** | *Asticcacaulis biprosthecum* | Kind gift from Yves Brun, UDEM |
| **GL2078** | *Asticaccaulis benevestitus* | Kind gift from Yves Brun, UDEM |
| **GL2079** | *Brevundimonas subvibriodes* | Kind gift from Yves Brun, UDEM |
| **GL2080** | *Phenylobacterium lituiforme* | Kind gift from Yves Brun, UDEM |
| **GL2296** | *Hyphomonas neptunium* | Kind gift from Yves Brun, UDEM |
